# T-Rex: sTandalone Recorder of EXperiments; An easy and versatile neural recording platform

**DOI:** 10.1101/2022.10.26.513822

**Authors:** Joaquín Amigó-Vega, Maarten C. Ottenhoff, Maxime Verwoert, Pieter Kubben, Christian Herff

## Abstract

Recording time in invasive neuroscientific empirical research is short and must be used as efficiently as possible. Time is often lost due to long setup times and errors by the researcher. Minimizing the number of manual actions reduces both and can be achieved by automating as much as possible. Importantly, automation should not reduce the flexibility of the system. Currently, recording setups are either custom-made by the researchers or provided as a module in comprehensive neuroscientific toolboxes, and no platforms exist focused explicitly on recording. Therefore, we developed a lightweight, flexible, platform- and measurement-independent recording system that can start and record experiments with a single press of a button. Data synchronization and recording are based on Lab Streaming Layer to ensure that all major programming languages and toolboxes can be used to develop and execute experiments. We have minimized the user restrictions as much as possible and imposed only two requirements on the experiment: The experiment should include a Lab Streaming Layer stream, and it should be able to run from a command line call. Further, we provided an easy-to-use interface that can be adjusted to specific measurement modalities, amplifiers, and participants. The presented system provides a new way of setting up and recording experiments for researchers and participants. Because of the automation and easy-to-use interface, the participant could even start and stop experiments by themselves, thus potentially providing data without the experimenter’s presence.

## 1 Introduction

Measuring high-quality electrophysiological human brain activity in neuroscientific research is notoriously difficult. High-quality signals can for example be measured from invasive electrodes, providing high spatial and temporal resolution ([1], [2]). However, since implanting humans solely for neuroscientific research is ethically debatable, researchers often ’piggy-back’ on clinical treatments of patients that receive electrode implants ([3], [4]). For example in patients with medication-resistant epilepsy undergoing presurgical monitoring for resection surgery ([5]) or patients qualified for deep brain stimulation ([6]).

The time to record neuroscientific experiments with these patient groups is severely limited. For example, epilepsy patients are usually in a monitoring unit ranging from a few days to two weeks, and for patients with deep brain stimulation, measurement windows are either microelectrode recordings during surgery or the week after surgery up until the stimulator is turned on. Measurements must not interfere with the clinical treatment, and patients need time to sufficiently recover to be able to participate, further reducing the available time. It is also common that several groups want to perform their own measurements, resulting in a further division on the already limited time.

For this reason, it is essential that the brief time window that remains is used as efficiently as possible. This means the amount of time spent on recording should be maximized, and inversely the time spent on setup and solving errors should be minimized. Both setup time and the error rate can be significantly reduced by automating as many manual actions as possible (e.g., connecting to recording devices, starting experiments, selecting data streams, starting, stopping, and synchronizing the recording). However, as experiments or recording setups change over time, it is often not worthwhile for research groups to invest in developing a more sophisticated system. It takes human resources, technical knowledge, and notable time investment to move beyond the prototypical system, which is unavailable to most research labs.

Aside from the custom made setups, there are a few widely used measurement platforms, such as BCI2000 ([7]), OpenVIBE ([8]) and FieldTrip ([9]). These systems can record data from a large number of different amplifiers, as well as provide modules to design and analyze experiments. When considering only the recording functionality, these platforms do not all run on all operating systems, are restrictive in either hard- or software toolset, programming language, input or output, or are not open-source. For example, FieldTrip requires the researcher to use the proprietary platform MATLAB, and the BCI2000 and OpenVIBE require the researcher to use their tools and API. Additionally, the user must install the full software package on their system, even when only the recording functionality is needed. While we think these platforms are valuable in their own right, we identified that a standalone platform is missing solely focused on recording data. Such a system should be able to run on any operating system, should not restrict the users to a specific set of hard- or software, and should be open-source. Ashmaig et al. ([10]) developed and described a system exclusively focused on continuous data recording for neurosurgical patients. The system provides a good use case for naturalistic long-term recordings but has an extensive list of hardware requirements and limits the user to Linux. Additionally, not all research groups have the opportunity to perform long-term recordings.

Therefore, we developed sTandalone Recorder of EXperiments or *T-Rex* for short, purposely designed for easy setup and recording of neuroscientific experiments. We ensured *T-Rex* is lightweight, open-source, and independent of toolset or operating system. The system can be used in any lab with access to a laptop and provides the researcher with an easy-to-use user interface. Moreover, we automate all setups and the start and stop of experiments, allowing the researcher to start experiments with a single press of a button.

## 2 T-Rex platform

As there are many differences between recording setup between labs, we set three criteria to make *T-Rex* applicable in most labs: first, *T-Rex* should be **independent** of the choice of tools, paradigms, operating systems and programming languages. Each lab has its preferred tool sets, and not restricting specific sets allows labs to port experiments into *T-Rex* easily. Secondly, *T-Rex* should be **user-friendly** for both researcher and participant. By increasing simplicity, error rates and time spent on setup can be reduced. Thirdly, the system should be **robust**, meaning that the experiment should always run, and in case of technical problems, retain the data up to that point and return to the *Home* screen.

### 2.1 System outline

In short, *T-Rex* acts as the middle man handling the experimental overhead for the researcher (Figure 1). When using *T-Rex*, the researcher can select an experiment by simply pressing a button on the main menu screen (Figure 2A). When an experiment is selected, *T-Rex* checks the availability of all required input devices for the selected experiment and establishes a connection. For example, a movement-tracking experiment requires input from a hand-tracking device and the amplifier measuring the participants’ neural activity. *T-Rex* will search for and connect to these devices. *T-Rex* will also start the experiment user interface (UI) that instructs the participant on what task to perform. It starts recording all data streams and saves them to a folder specified by the researcher. The synchronized data is saved inside this folder using the `.xdf` file format. Lastly, the UI prompts the participant on how the experiment went and returns to the *Home* screen, where the next experiment can be selected. The only action the researcher has to perform is to start the required device data stream(s) and select the experiment in the *Home* screen.

**Figure 1:**
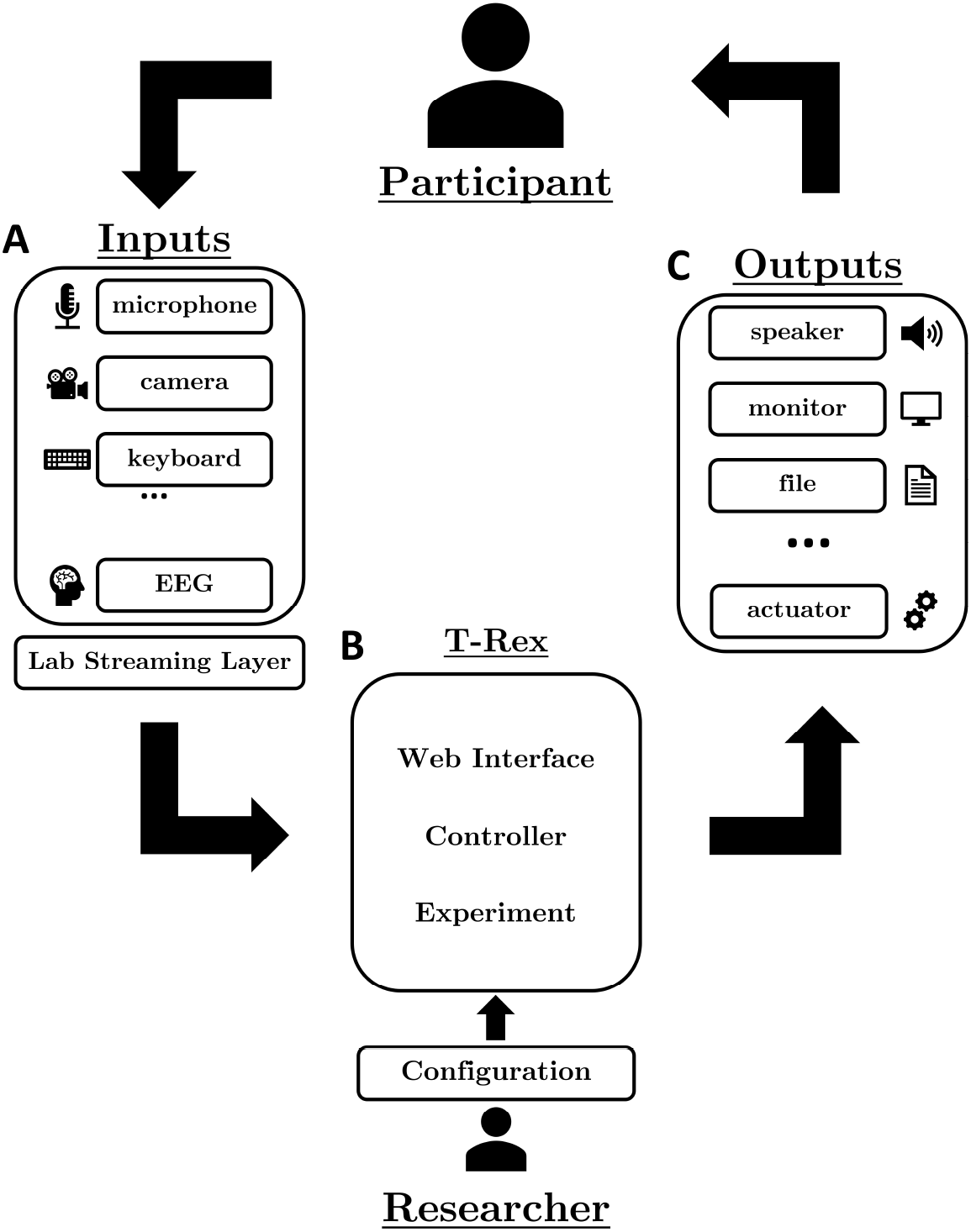
The use of *T-Rex* imposes no limitations on the inputs, outputs, behavior, or frameworks used by the researchers for creating the experiments. **(A)** Represents a set of possible inputs. **(B)***T-Rex* sits in the middle handling the logic that connects the inputs and the outputs. The three main software components of the system are also illustrated (Web interface, Controller, and User configuration). **(C)** Depicts a set of possible outputs. It is worth emphasizing that the inputs and outputs are not limited to those represented.

**Figure 2:**
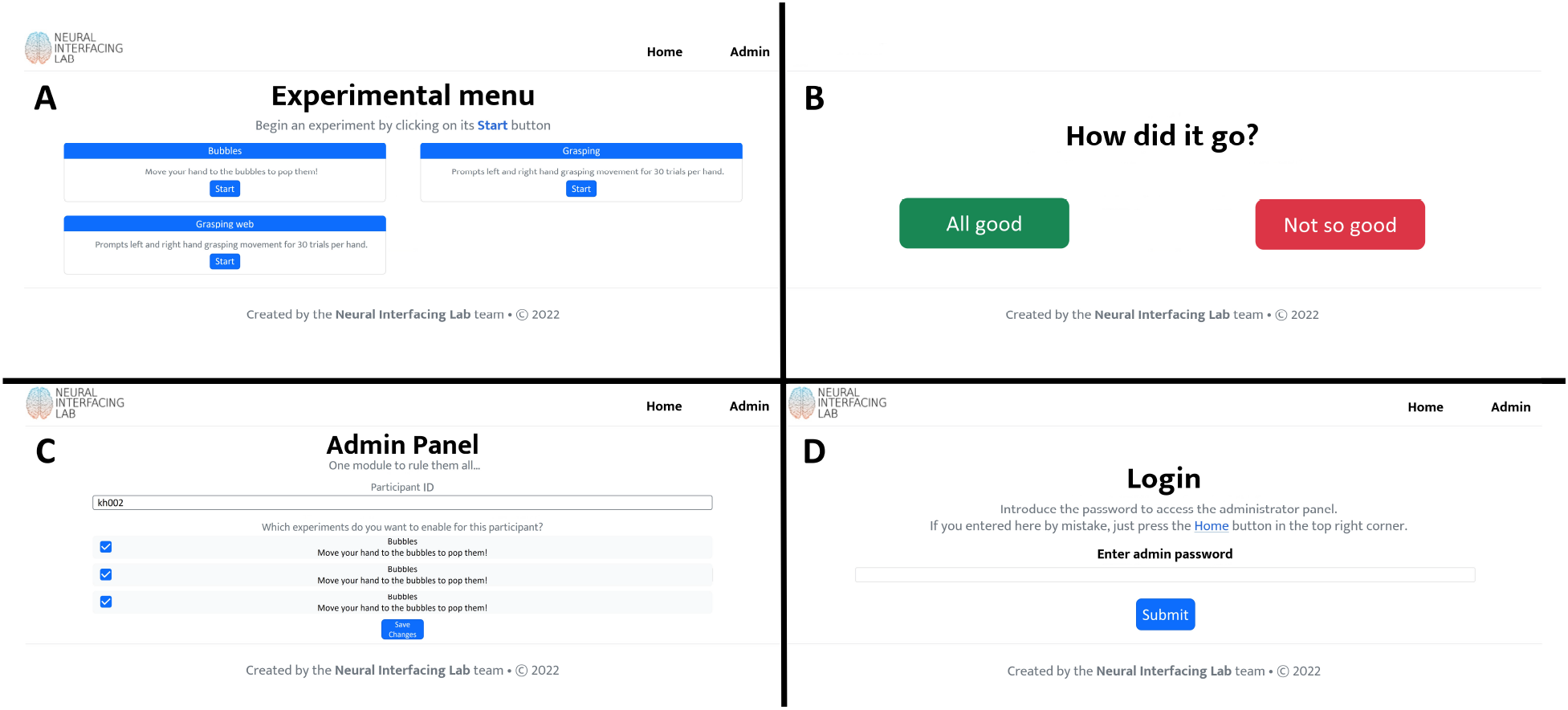
Representation of the main four windows of the Web interface. **(A)** The *Home* window contains all the experiments accessible to the user, represented on a grid configuration. **(B)** The *Experiment Feedback* window allows obtaining feedback from the participants about their experience with the experiment. It is achieved through the green **(“All good”)** and the red **(“Not so good”)** button. The participants can only continue after pressing one of these buttons. **(C)** The *Admin Login* window allows access to the administration panel by entering the password. **(D)** The *Admin Configuration* window, where the administrator can create new participants and modify their access to experiments.

#### 2.1.1 Materials, software, and technologies

The web interface (including the *Home* screen with experiment selection, see Figure 2A-D) is built using Bootstrap5^1^ for the front-end and the Python package Flask^2^ for the back-end. The Controller, handling set up, start and stop of the experiments (section 2.2.2), is built in Python 3.9+ and requires a few dependencies found in `requirements.txt` in the repository. Data synchronization is implemented using Lab Streaming Layer ([11]).

*T-Rex* is compatible with Windows, macOS, and Linux. As *T-Rex* is lightweight and primarily active before and after experiments, the hardware requirements are determined mainly by the recording of the experiment in Lab Streaming Layer, instead of *T-Rex*.

##### Lab streaming layer

*T-Rex* uses Lab Streaming Layer (LSL) to synchronize the data streams from different devices, such as a variety of EEG amplifiers, audio streams, movement trackers, and cameras. The service handles *“networking, time-synchronization, (near-) real-time access and optionally the centralized collection and recording of data.”* ^3^. It is lightweight and has multi-language and platform support, including Unity and Android. LSL allows the user to send data via a data stream to a local network server which can be recorded. Basic usage involves instantiating a StreamOutlet object with a name, type, channel count, sample rate, data type, and source id. For example, the most basic application in Python is represented in the following code-block:

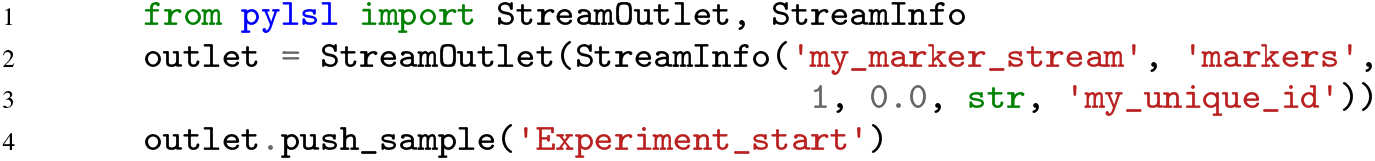

Inversely, to receive data one can instantiate a `StreamInlet` and use `inlet.pull_sample()`. For a comprehensive overview, see the official documentation.^4^ *T-Rex* requires each device or experiment included to start a `StreamOutlet` similar to the example above. If no `StreamOutlet` is created, *T-Rex* will not be able to find the device and start the experiment.

##### Trigger

In some recording setups, a trigger is used to mark the start and end of an experiment. In these setups, the participant’s clinical data is recorded continuously and stored on a server. During an experiment, the data cannot be streamed directly and needs to be retrieved afterwards by the responsible data steward. The data steward can locate the requested data files by sending a unique trigger pattern (usually represented as an on-off signal) directly to a separate channel in the amplifier. T-Rex includes specific functionality to send a trigger signal at the start and end of the experiment.

### 2.2 Software components

The software consists of two main components: the web interface that handles the UI and the Controller that sets up, starts, and stops all experiments (Figure 1B).

#### 2.2.1 Web interface

The Web interface includes four windows: *Home*, *Experiment Feedback*, *Admin Login*, and *Admin Configuration* (Figure 2).

The *Home* window (Figure 2A) displays all the experiments in a grid. Experiments’ cards’ are shown on that grid with a title, description, and start button. When the button is pressed, the *Controller* executes a command that starts the selected experiment. The command is defined by the researcher and specified on the configuration of the experiment (more details in section 2.3). During the experiment, the web interface is on stand-by awaiting the completion of the experiment.

After completion, the participant is redirected to the *Experiment Feedback* window where the question *“How did the experiment go?”* is prompted (Figure 2B). The participants are required to select a feedback option to continue. This allows the researcher to save a brief experiment evaluation to assess data quality in later analysis. In potential future applications, the participant might perform the experiments by themselves. Then this feedback is useful to flag the researcher to be aware of a potential loss in data quality during later analysis. The feedback is stored under the file name feedback.txt in the same folder as the most recent `.xdf` file (that contains the data recorded from the experiment).

The *Admin Configuration* provides the researcher with a closed environment where the participant identifier and a selection of all available experiments can be chosen. To access the *Admin Configuration*, the researcher must first log in using the password that is configured in the main configuration file (Figure 2C, 2.3 for details). When logged in, the researcher can see the configuration of the active experimental session, composed of an alphanumeric participant identifier and their access to experiments. A list of all the experiments included in the platform is visible from this window, but only those with checked marks are visible to the participant. The changes on this window are only applied after pressing the **“Save”** button at the end of the page.

#### 2.2.2 Controller

The Controller handles everything related to running an experiment and has three main parts: 1) Setup, 2) Start, and 3) Stop (Figure 3). The related code can be found in the `./libs directory`.

**Figure 3:**
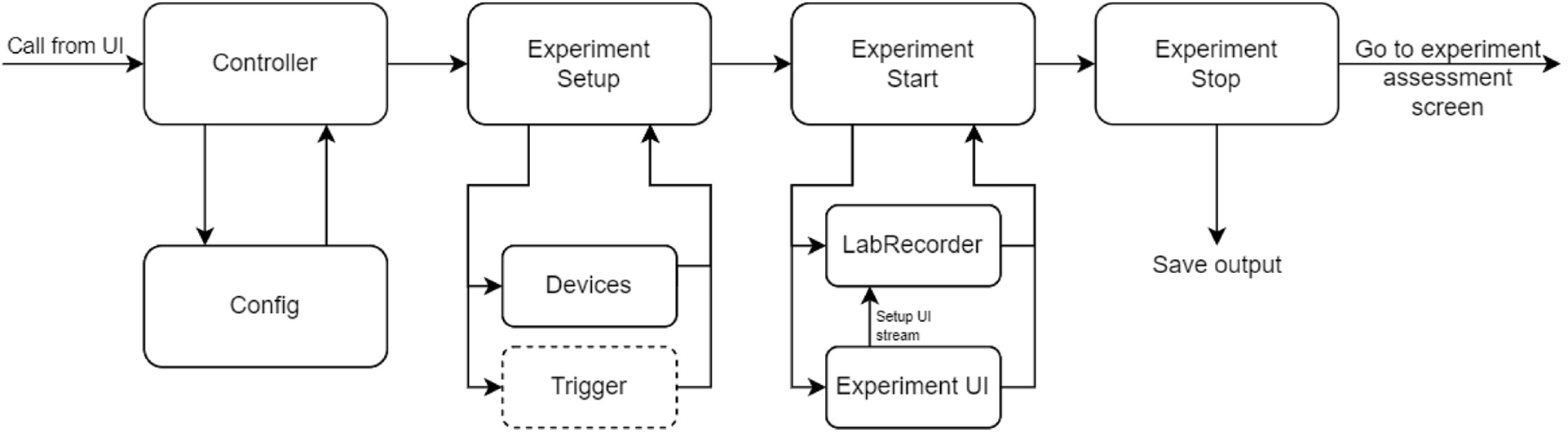
Flow of running an experiment. When an experiment is started by pressing the start button on its card, the `Controller` is called, loading the main configuration file and extracting the information received from the UI about which experiment to run. Then an `Experiment` instance is created, loading the experiment-specific information and completing the setup in three steps. First, it checks for all devices and their LSL streams. Second, it initializes a Recorder instance and adds all streams to the list of streams it should record. Third and last, if a trigger is required for the selected experiment, it will set up a trigger class that searches and connects to the trigger. Once the subprocess call is returned, `Experiment` sends the final trigger and stops the Recorder. The data is saved in the `./output/` folder, and the user is redirected to the experiment assessment screen (Figure 2B).

##### Setup

When an experiment is started by pressing the start button on its card, the `Controller` class in *Controller.py* (Figure 3) is called, loads the main configuration file and extracts the information received from the UI about which experiment to run. With this information, an `Experiment` instance is created, and its loading function is called.

`Experiment` loads the experiment-specific information and completes the setup in three steps. First, it checks for all devices and their LSL streams as defined by the user in the experiment configuration under device_inputs.

Subsequently, `Experiment` initializes a Recorder instance and adds all streams to the list of streams it should record. For a movement experiment ([12], [13], [14], [15], [16]), the streams recorded could be the neural amplifier, experimental triggers, and a movement tracker could be potentially added ([17], [18], [19]) or a force sensor ([20]. For speech perception ([21], [22], [23]) or auditory perception ([24], [25]) the audio stream, experiment triggers and neural data. For speech production ([26], [27], [28], [29]), the streams could be neural data, microphone, and triggers. In section 3 we provide some example experiments.

The last step is to check if a trigger is required for the selected experiment. If positive, it will set up a trigger class that searches and connects to the trigger.

It is essential that all devices are connected and active before the `Experiment` instance is called. As all requested devices are essential for successful recording, *T-Rex* will raise an error and return to the UI if not all input devices are connected successfully.

##### Start

A user-defined command is called using Python’s subprocess library to start the experiment UI. This command is set in the experiment-specific configuration and should be capable of being executed from the command line interface. Because the experiment UI likely contains a stream that sends out experiment-related markers, `Experiment` will start a loop on a user-defined timeout to search for the marker stream. Once found, usually, almost instantly, the Recorder will start recording all streams. Implementing the system this way does not restrict the research aside from using LSL. However, due to the timeout, the experiment may start before the recording starts. This can only happen if the time between the setup of the experiment StreamOutlet and sending of the first marker is shorter than the time that the Recorder can find the stream and start the recording. Usually, finding the StreamOutlet and start recording is in the order of milliseconds. However, to entirely prevent the possibility of this happening, we recommend including a waiting screen in the experiment UI (i.e., “Press button to start”) or built in sufficient time (longer than the timeout set in the experiment configuration) between the setup of a StreamOutlet and the start of the experiment. Once connected to the experiment StreamOutlet, the experiment UI should start and the `Experiment` instance will wait until the called command is terminated and returns, which usually happens when the experiment UI window is closed.

##### Stop

Once the subprocess call is returned, `Experiment` sends the final trigger and stops the Recorder. The data is saved in the `./output/` folder, defined in the main configuration file (section 2.3), an example of the created directory tree can be found in the Supplementary Figure S4.

#### 2.2.3 Device inputs

Each experiment can have multiple input devices, such as the amplifier measuring the neural data, hand-tracking devices or a microphone. Any device can be included as long as it generates a StreamOutlet. Each device should send the data from the device to LSL, allowing it to be accessed by the other system components and to be recorded. The name, type or source_id supplied to the StreamOutlet will be the values that *T-Rex* will search for at experiment setup (section 2.2.2). In practice, this means that either the name, type, or source_id needs to be supplied under device_inputs in the experiment configuration file (section 2.3.2). Since devices can be used for multiple experiments, we included a separate destination for all device input files: `./exp_module/inputs`, although input devices can be stored anywhere as long as they generate a StreamOutlet.

### 2.3 User configuration

There are two types of configuration files that the user can set: the main configuration and experiment-specific configurations. All configuration files are formatted in Yet Another Markup Language (YAML).

#### 2.3.1 Main configuration

The file `config.yaml` on the root folder contains the system-wide configurations. This configuration file contains information containing general settings. Supplementary Figure S1 contains a description of the different available options and Supplementary Figure S2 contains an example of the main configuration file. The `main` option under `path` is the path all relative paths will be anchored to and should be set to the root folder. Most parameters are preset, but out and `trigger` configurations may vary between different measurement setups and might need to be redefined.

#### 2.3.2 Experiment configuration

Each experiment included in *T-Rex* requires a separate folder in `./exp_module/experiments/` and must include at least two files: `config.yaml` and the entry file to start the experiment. In this experiment configuration file, `name` and `description` define the text shown in the UI. `command` sets the command line interface command made by the controller class to start the experiment. `exp_outlet` sets the name, type or source_id that the Experiment class (section2.2.2) will search for, for `timeout` seconds. For example, if your experiment UI is a Python script that will create a StreamOutlet named *markers*, then `command` would be python `.\exp_module\experiments\your_experiment_file.py and exp_outlet=`’markers’. A full description of all experiment configuration fields and options can be found in Supplementary Figure S5.

## 3 Use cases

We have included three different example experiments to provide a more practical view of how to use *T-Rex*. The examples can also serve as a quick start for researchers to create new experiments or adapt the ones included. A step-by-step explanation of adding a new experiment is described in section 3.4.

### 3.1 Case 1: Simple experiment in Python

This experiment is a simple text-based instruction for a grasping task (Figure 4A). The participant is prompted by text in a Python tkinter^5^ window to continuously open and close either their left or right hand, as used in [13]. The experiment requires neural data as the input device and generates a StreamOutlet to send markers that inform about the start and end of the experiment and of the trials. The neural data is acquired from a stream with *name* = *Micromed*, *type* = *EEG*, *source*_*id* = *micm*01. These values are all set by the user. As *T-Rex* will search for all three options (name, type or source_id), only one needs to be provided. Therefore, the option under `device\_inputs` in `grasping\config.yaml` is set to `eeg` (case-insensitive). Next, the Marker StreamOutlet that the experiment itself will generate has *source*_*id* = *emuidw*22. When the `Experiment` class runs the experiment command (`command` field in `grasping\config.yaml`) it will search for this streams. Therefore, the `exp_outlet` field is set to ’emuidw22’. Finally, since the grasping experiment is Python-based, the `command` should use Python to call the script with the command: `python .\exp_module\experiments\grasping\grasping.py`. The configuration file used can be found in Supplementary Figure S6.

**Figure 4:**
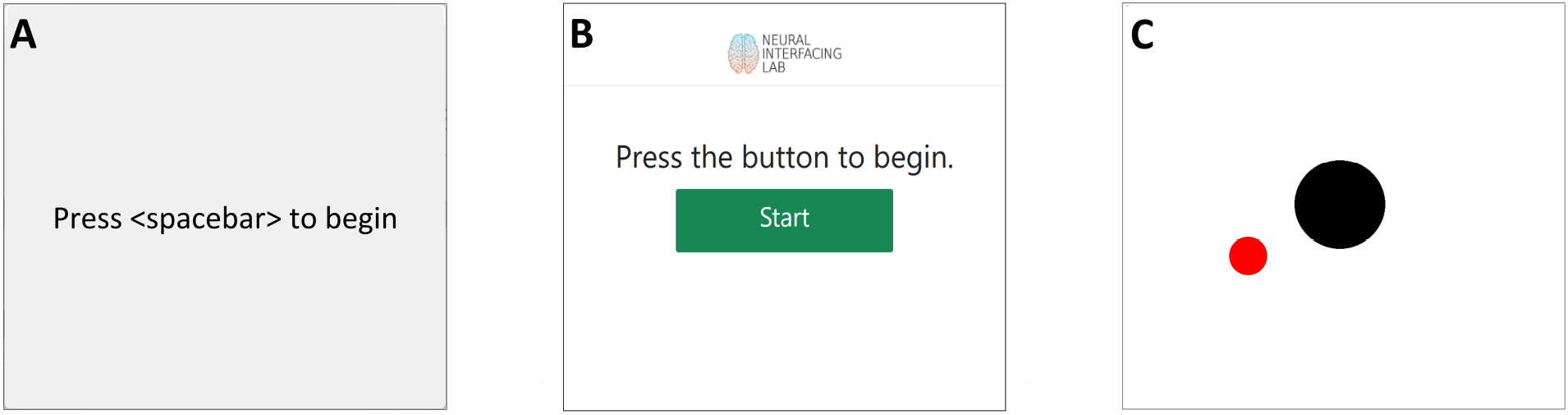
User interfaces for the three Use cases experiments included. **(A)** Grasping, simple text-based experiment built using the Python package Tkinter (Tk). **(B)** Grasping Web experiment, re-implementation of the Grasping experiment as a Single Page Application (SPA) to allow its execution on any device with access to a web browser. **(C)** 3D hand-tracking experiment, the hand-tracking is performed using the LeapMotion controller, and the experiment is implemented in Python using the package Tkinter (Tk).

When these options are set, the experiment is ready to go and can be started by pressing the start button on the *Home* window. The Tk window opens and waits for the spacebar to be pressed. Once pressed, the experiment starts and is locked on the topmost position upon completion. When the experiment is finished and closed (i.e., the command call ends and returns to the Experiment class), the `Experiment` instance stops the recording and saves the data. In-depth details on how experiments are started and stopped are described in section 2.2.2.

### 3.2 Case 2: Simple experiment in a WebUI

From the Web Interface, both participants and researchers can access the main functionalities of *T-Rex*, allowing the potential execution of the system on a headless server, different from the interface that interacts with the participants. The system that runs *T-Rex* could create a local network that serves the experiments to different devices (tablets, laptops, smartphones), removing the need for an active internet connection which could render the execution unsafe.

To illustrate this paradigm, we created the grasping web experiment, which mimics the behaviour of the “grasping” experiment (section 3.1) but in a web format (Figure 4B). Given that this experiment consists of a Single Page Application (SPA), it can be wholly executed on any device with access to a web browser, like laptops, tablets, and smartphones. The grasping web experiment also illustrates options other than a Tk window for experimenting.

For constructing the experiment, we used HTML, CSS (using Bootstrap5 for the responsiveness and other visual aspects), and JavaScript for the behavior. The device input is the same as in the Tk implementation as well as the StreamOutlet containing the markers, thus the `device_inputs` and `exp_outlet` are the same. The difference is in the command executed to start the experiment. In this case, `start .\exp_module\experiments\graspingWeb\index.html` is used. The configuration file used can be found in Supplementary Figure S7.

Once the experiment is started on the *Home* window, the `Experiment` instance opens another tab on the browser displaying the “grasping_web” experiment. The experiment starts when the participant presses the green “Start” button. When the experiment is finished, the participant or researcher is prompted to press a red button to close the experiment. The GraspingWeb command call is finished at button-press and returns to the `Experiment` instance, stopping the recording and saving the data.

### 3.3 Case 3: Multiple devices

Lastly, we included a 3D hand-tracking experiment, where the goal is to hold the cursor (a black circle) on the target (a red circle). The cursor can be moved in 3d, where the third dimension controls the size of the circle (Figure 4C). In this case, the hand-tracking is done by the LeapMotion controller^6^. We have provided a .exe that reads the data from the tracker and sends it to an LSL StreamOutlet with *name* = *LeapLSL*, *type* = *Coordinates*, *source*_*id* = *LEAPLSL*01. In addition to the hand-tracking information, we also need neural activity, for which we use the same StreamOutlet as described in case 2 (section 3.1). Lastly, the experiment is implemented in a Python Tk window and generates a marker Stream similar to the streams described in the previous use case with *Source*_*id* = *BUBBLE*01. Thus, to set up the configuration for this experiment, we set command to `python .\exp_module\experiments\Bubbles\bubbles.py`, exp_outlet to `BUBBLE01` and device_inputs to `LEAPLSL01` (the tracking information stream) and eeg (the neural data stream). To run the experiment, the researcher should start the device stream before the experiment is started in the *Home* screen. (i.e. run the `.exe` first). The configuration file used can be found in Supplementary Figure S8.

#### 3.3.1 Mix ’n Match

These are just three examples showing different possibilities. It is easy to include more devices by simply adding a StreamOutlet name, type, or source_id to the list of `device\_outputs`. However, it is required that the researcher writes a script to read the data from the device and send it to an LSL StreamOutlet. New experiments can also be built in Unity^7^ or PyGame^8^ to provide better graphical experiences. As mentioned before, we do not impose restrictions on which technologies to use or the specific type of experiments that can be performed, whether speech production, audio or speech perception, movement, or simple or more naturalistic tasks ([30], [31]).

### 3.4 Adding new experiments to the platform

The following steps describe how to add a new experiment from scratch to *T-Rex*:

1. Create the experiment folder inside the directory `./exp_module/experiments/`. An example of the directory tree for different example experiments can be found in Supplementary Figure S3.
2. Create the experiment configuration file (`config.yaml`) inside the new folder. Supplementary Figure S9 can be used as the base example for creating this file, and section 2.3.2 contains a detailed description of each parameter.
3. Adjust the fields to the specific experiment (section 2.3).

After completing these initial steps, the experiment should be visible from the *Admin Configuration* panel. The researcher can set the experiment as “visible” from the admin panel by selecting its corresponding check mark. If configured as “visible”, it should appear on the *Home* window, and it can be executed by clicking on its respective button.

It is worth mentioning that when porting an already-configured version of *T-Rex* to a different OS, some parameters might need to be revised. For example, the parameter `command`, when used on Windows to start a python experiment, the definition is the following:

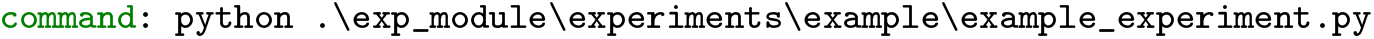

However, when used on Unix or Unix-like systems, the definition changes to the following:

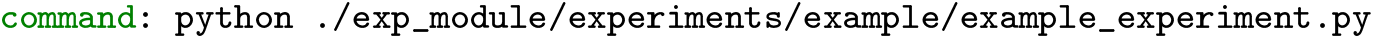

The difference comes because “/” is the path separator on Unix and Unix-like systems, and Microsoft uses “\”.

There might be other scenarios where the parameter `command` might differ between OS, so we recommend revising each experiment configuration file when porting the platform to a different OS.

Even so, incorporating new experiments into *T-Rex* is seamless. Several experiments can be quickly integrated into the platform with just a few easy steps.

## 4 Discussion

We presented *T-Rex*, an independent, user-friendly, and robust system that minimizes setup time and error rate. *T-Rex* provides a simple UI and reduces the experimental setup to a press of a button. The software merges features for automated continuous recordings and synchronization of neural data into an easily operated web interface. Once coupled to *T-Rex*, the experiments can be executed directly from the interface. It also includes functionalities for managing several participants with different access levels.

The simplicity of *T-Rex* reduces the amount actions that the researcher has to make to two: Starting the required device(s) and starting the experiment. This significantly reduces the room for error, saves time to set up and check all connections, improves reliability, and leaves more time for actual experiments. We aimed to make *T-Rex* as flexible as possible by making it available to all operating systems, most main programming languages (limited by LSL), and web browsers. So far, we can verify that it works with Firefox (version: 105.0.1), Chrome (version: 106), Safari (version: 16), and Edge (version: 106), although it should be compatible with higher versions and other mainstream browsers. Any software that provides a command line interface is compatible with *T-Rex* (for example, Psychopy^9^, PyGame, Unity). This also means that a system can easily be transferred to other devices by copying the root folder to the new device and installing the required dependencies. It is noteworthy that measurements with *T-Rex* are not limited to neuroscientific research. Lastly, as we used LSL to handle time series synchronization, there are also networking options, allowing for data recording over a network of devices instead connections to a single device.

Relying on a command line interface command to start an experiment also means that it does involve some knowledge of shell scripts. The commands or scripts are platform dependent and will likely need to be adjusted to the OS where the platform needs to be executed, even when executing the same experiments. Additionally, experiments need to include a StreamOutlet object in the code to be included in *T-Rex*, which requires some adjustment to existing experiments and programming knowledge. The same is true for the device inputs required for experiments. This usually means the researcher needs to create a basic script that reads data from the device and sends it to LSL. The researcher should also start these device scripts before starting the experiment.

*T-Rex* is in ongoing development, and starting the device scripts at the experiment start is likely to be added in future updates. We have designed *T-Rex* so that in future updates, it is not necessary anymore for the researcher to be present during experiments. If experiments are engaging enough, the participants may want to start an experiment themselves. *T-Rex* contains the core structure to automate the recording process entirely, which could increase the amount of experimental neural data gathered.

To summarize, we developed a new recording platform that is Operating System independent, user-friendly and robust. We provide researchers with a solution that can significantly increase time spent on recording instead of on the setup with its possible errors.

## Code availability

The source code, installation guide, and example experiments can be found on GitHub ^10^. *T-Rex* is available under the permissive MIT License. As *T-Rex* will be in ongoing development, we kindly invite users to provide feedback or contribute to this open-source project.

## Conflict of Interest Statement

The authors declare that the research was conducted in the absence of any commercial or financial relationships that could be considered as a potential conflict of interest.

## Author Contributions

*Conceptualization*: JAV, MCO, MV, PK, CH *Data Curation*: N/A *Formal analysis*: N/A *Funding acquisition*: N/A *Investigation*: JAV, MCO, PK *Methodology*: JAV, MCO, PK, CH *Project administration*: JAV, MCO *Resources*: PK, CH *Software*: JAV, MCO, PK *Supervision*: PK, CH *Validation*: JAV, MCO, MV *Visualization*: JAV, MCO *Writing—Original Draft*: JAV, MCO *Writing—Review & editing*: JAV, MCO, MV, PK, CH

## Funding

CH acknowledges funding by the Dutch Research Council (NWO) through the research project’ Decoding Speech In SEEG (DESIS)’ with project number VI.Veni.194.021.

## Abbreviations

T-Rex: sTandalone Recorder of EXperiments
UI: User interface
CLI: Command Line Interface
OS: Operating System
YAML: Yet Another Markup Language

## Footnotes

1 https://getbootstrap.com/docs/5.1/getting-started/introduction/

2 https://flask.palletsprojects.com/en/2.1.x/

3 https://github.com/sccn/labstreaminglayer

4 https://labstreaminglayer.readthedocs.io/info/getting_started.html

5 https://docs.python.org/3/library/tkinter.html

6 https://www.ultraleap.com/product/leap-motion-controller/

7 https://unity.com/

8 https://www.pygame.org/wiki/about

9 https://www.psychopy.org/

10 https://github.com/neuralinterfacinglab/t-rex

## Notes

https://github.com/neuralinterfacinglab/t-rex

### Competing Interest Statement

The authors have declared no competing interest.

