## Supplementary Document for "T-Rex: sTandalone Recorder of EXperiments; An easy and versatile neural recording platform"

```
path:
  main: Path to the root directory of the project, when set to
        "default" the value is obtained automatically.
  exe:  Relative path to the Lab Recorder executable.
  out:  Relative path to the output directory where the files
        generated during the experiments will be saved.

log_file: Relative path to the log file.
password: Password to access the Admin Configuration window.
debug:    Used for debugging purposes, when "True" a testing
          LSL stream under the name DBG01 will be created

trigger:
  sequence: Binary sequence that the trigger will send to
            the amplifier.
  serial_com_name: Name of the trigger device that the
                   recorder will search for.
```

**Fig. S1.** This is the system-wide configuration file that must be placed inside the root folder of the project, which allows the researcher to configure the execution of T-Rex.

```yaml
path:
  main: default
  exe: libs/labrecorder/LabRecorderCLI.exe
  out: output
log_file: record.log
password: 'qwer'
debug: False
trigger:
  sequence: [1, 0, 0, 1, 0, 0, 1]
  serial_com_name: 'Prolific PL2303GT'
```

**Fig. S2.** An example of a main configuration file example. Note that all paths are relative to the `main` parameter.

```
exp_module
├── inputs
└── experiments
    ├── EXPERIMENT_1
    │   ├── config.yaml
    │   └── experiment_1.py
    ├── EXPERIMENT_2
    │   ├── config.yaml
    │   └── experiment_2.html
    └── EXPERIMENT_3
        ├── config.yaml
        └── experiment_3.exe
```

**Fig. S3.** The directory tree illustrates a system with 3 different folders each for a different experiment ( ∼/EXPERIMENT_1/ , ∼/EXPERIMENT_2/ and ∼/EXPERIMENT_3/). Each experiment contain their own configuration file (config.yaml). The researcher can add any additional files to each folder.

```
output
└── PARTICIPANT_ID
    ├── access.yaml
    └── YYYY_MM_DD
        └── ID_OF_EXPERIMENT_HH_MM_SS
            ├── output.xdf
            ├── feedback.txt
            └── logs.log
```

**Fig. S4.** The directory tree illustrates the content of the `./output/` folder when saving the experimental data gathered with one experiment. The `output.xdf` file is created upon the experiment completion. It contains the recorded data from the pre-configured LSL streams. The `feedback.txt` file contains the feedback the participant inputted on the *Experiment Feedback* window and it is saved in the same folder as the most recent `.xdf` file.

```
name: Name of the experiment. This value will be used as the title for
      the web interface buttons shown to the participants.
description: Short description of the experiment that will be shown
             in the experiment "cards" in the user UI.
id: Unique identifier for the experiment. This name should match the
    name of the folder where the experiment is stored inside the
    directory "./exp_module/experiments".
command:  Command that is called by the Experiment instance. It must be able
          to run from a command prompt.
timeout: Amount of seconds the recorder will search for a marker stream.
exp_outlet: Source id of the marker stream generated by
            the user experiment (by calling "command").
device_inputs: List of devices that the recorder will search for.
               The recorder will find matching streams based on name,
               type, and source_id as defined by LabStreamingLayer.
trigger: Switch whether a trigger should be sent for this experiment.
```

**Fig. S5.** The different options for the experiment configuration file. Each experiment must include this file. The parameter `command` might need to be modified when porting the platform to a different Operating System (from Windows to Linux or macOS, for example). It is up to the researcher to perform the redefinition.

```
name: 'Grasping'
description: 'Prompts left and right hand grasping movement for
    30 trials per hand.'
id: grasping
command: python .\exp_module\experiments\grasping\grasping.py
timeout: 5
exp_outlet: emuidw22
device_inputs:
  - EEG
trigger: False
```

**Fig. S6.** Experiment configuration file used for the grasping experiment. This experiment presents simple instructions to the participant indicating continuous opening and closing of either their left or right hand. The visual interface was built using the Python-TK library.

```yaml
name: 'Grasping Web'
description: 'Prompts left and right hand grasping movement for
    30 trials per hand (using a web interface).'
id: graspingWeb
command: start .\exp_module\experiments\graspingWeb\index.html
timeout: 10
exp_outlet: GRASPW01
device_inputs:
  - EEG
trigger: False
```

**Fig. S7.** Experiment configuration file used for the grasping-web experiment. This experiment presents simple instructions to the participant indicating continuous opening and closing of either their left or right hand. The visual interface was built using HTML, CSS (Bootstrap5 for the responsiveness and other visual aspects), and JavaScript for the behavior.

```yaml
name: 'Bubbles'
description: 'Move your hand to the bubbles to pop them!'
id: bubbles
command: python .\exp_module\experiments\Bubbles\bubbles.py
timeout: 10
exp_outlet: BUBBLE01
device_inputs:
  - LEAPLSL01
  - EEG
trigger: False
```

**Fig. S8.** Experiment configuration file used for the 3D hand tracking experiment. The goal of the experiment is to hold the cursor on the target. The cursor can be moved in 3d, where the third dimension controls the size of the circle. In this case, the hand tracking is done by the LeapMotion controller.

```yaml
name: 'NAME OF THE EXPERIMENT'
description: 'DESCRIPTION OF THE EXPERIMENT'
id: 'ID OF THE EXPERIMENT, IT MUST MATCH THE FOLDER NAME'
command: 'COMMAND THAT EXECUTES THE EXPERIMENT'
timeout: 'TIMEOUT FOR STOP SEARCHING THE STREAMS (SECONDS)'
exp_outlet: 'ID OF STREAM GENERATED WITH COMMAND'
device_inputs:
    - 'DEVICE THE RECORDER WILL SEARCH'
    - 'DEVICE THE RECORDER WILL SEARCH'
trigger: 'True FOR USING TRIGGER, False OTHERWISE'
```

**Fig. S9.** Template that can be used for creating some experiment configuration file.
